# The gut microbial signature of gestational diabetes mellitus and the association with diet intervention

**DOI:** 10.1101/2021.09.07.459364

**Authors:** Na Wu, Jingwei Zhou, Heng Mo, Qing Mu, Huiting Su, Mei Li, Yimeng Yu, Aiyu Liu, Qi Zhang, Jun Xu, Weidong Yu, Peng Liu, Guoli Liu

**Affiliations:** Department of Central Laboratory & Institute of Clinical Molecular Biology, Peking University People’s Hospital, Beijing, China, 100044; Department of Gynecology and Obstetrics, Peking University People’s Hospital, Beijing, China, 100044; Department of Stomatology, Peking University People’s Hospital, Beijing, China, 100044; Department of Clinical Nutrition, Peking University People’s Hospital, Beijing, China, 100044

## Abstract

Gestational diabetes mellitus (GDM) is a high-risk pregnancy complication that is associated with metabolic disorder phenotypes, such as abnormal blood glucose and obesity. The link between microbiota and diet management contributes to metabolic homeostasis in GDM. Therefore, it is crucial to understand the structure of the gut microbiota in GDM and to explore the effect of dietary management on the microbiota structure. In this study, we analyzed the composition of the gut microbiota between 27 GDM and 30 healthy subjects at two time points using Illumina HiSeq 2500 platform. The taxonomy analyses suggested that the overall bacteria clustered by diabetes status, rather than diet intervention. Of particular interest, the phylum *Acidobacteria* in GDM was significantly increased, and positively correlated with blood glucose levels. Moreover, Partial least-squares discriminant analysis (PLS-DA) revealed that certain genera in the phyla *Firmicutes, Bacteroidetes, Proteobacteria*, and *Lentisphaerae* characterized the GDM gut microbiota. Correlation analysis indicated that blood glucose levels and BMI index were correlated with the relative abundance of SCFAS-producing genera. Through the comparison between the GDM and healthy samples with or without diet intervention, we discovered that the role of short-term diet management in GDM processes is associated with the change in the *Firmicutes/Bacteroidetes* ratio and some specific taxa, rather than an alternative gut microbial pattern. Our study have important implications for understanding the beneficial effects of diet intervention on the specific gut microbiota and thus possibly their metabolism in pregnant women with GDM.

**Importance:** Understanding the composition and dynamics of the gut microbiota in GDM women under diet intervention is important because there may be opportunities for preventive strategies. We examined the relationships between GDM gut microbiota at two times before and after the diet intervention during second trimester of pregnancy and clinical characteristics in cohort of GDM women. We found that short-term diet management in GDM processes is associated with changes in the *Firmicutes/Bacteroidetes* ratio and some specific taxa rather than an alternative gut microbial pattern. Our study highlights the importance of considering diet intervention as the rescue of microbial dysfunction of GDM disease and can serve as a strategy for early prevention in future study.

## Introduction

The intestinal microbiota is a robust ecosystem inhabited by nearly 100 trillion bacteria (1). In recent years, extensive attention has been given to the gut microbiota during pregnancy. Over the course of a healthy pregnancy, the body undergoes substantial hormonal, immunological, and metabolic changes (2, 3). In predisposed women, these physiological changes may lead to the development of gestational diabetes mellitus (GDM). GDM is defined as abnormal glucose regulation with onset or first recognition during pregnancy and is one of the most common complications during pregnancy, with an incidence of 2–6% of all pregnancies (4, 5). The clinical incidence of GDM in China is currently presenting a dramatic increasing trend (6). In the context of nonpregnant obesity, recent work suggests a role for gut microbiota in driving metabolic diseases, including diabetes, weight gain, and reduced insulin sensitivity (4, 5, 7, 8). Researchers understand that the intestinal flora has an important function in the development of GDM with the notions relating the intestinal flora to metabolic disease (3, 9, 10). GDM is a transient state, and GDM patients are commonly treated by diet management to keep blood glucose within the normal range and reduce the risk of GDM complications (11). However, very few data from observational studies are available about whether diet interventions performed on GDM patients affect the community structure of the gut microbiota. Diet, particularly long-term eating habits, is known to be one of the drivers of microbiota variation (12, 13). Recent clinical studies have shown the importance of routine dietary recommendations for GDM patients, showing a better microbial pattern at the end of the study (14). However, the comparison between healthy pregnant women without dietary recommendations and individuals with GDM under routine dietary management remains uncertain.

In this study, we characterized the different patterns of the gut microbiota between GDM and healthy pregnancies in the second trimester of pregnancy. Then, comparison of microbial structure between healthy pregnant women without dietary recommendations and individuals with GDM under routine dietary management were assessed, to evaluate the role of short-term diet management on GDM gut microbiota. The aim of the present study was to provide an update on the existing knowledge of the specific structure of the gut microbiota in Chinese GDM women and to elucidate the influence of diet management on the GDM gut microbiota.

## Material and methods

### Patient recruitment

This study was approved by the Conjoint Health Research Ethics Board of Peking University People’s Hospital, and informed consent forms were signed by all of the subjects prior to participation in this study. All experiments were performed in accordance with the approved guidelines and regulations.

Diagnosis of GDM is based on the results of the fasting 75 g OGTT at 24–28 weeks gestation. One or more elevated level(s) is sufficient for a diagnosis of GDM. The threshold values of OGTT (5.1 at 0 hour, 10.0 at 1 hour and 8.5 at 2 hours during OGTT) are based on the diagnostic criteria recommended by the International Association of the Diabetes and Pregnancy Study Groups in 2011.

Thirty healthy subjects were selected based on matched age and pregnancy period, no complicating diseases and no antibiotic use during the 3-month period prior to sample collection. All subjects who met the following criteria were excluded: complicating diseases (such as known diabetes mellitus, hypertension, cardiovascular, pulmonary, autoimmune, joint, liver or kidney diseases; thyroid dysfunction; or any other disease), prebiotics/probiotics use, and antibiotic use during pregnancy.

The prepregnancy weight was self-reported; weight and height were measured at the time of enrollment. BMI was calculated as weight divided by the square of height. Arterial blood pressure (BP) was measured from the left arm with the participant in a sitting position after at least 10 min of rest with a mercury sphygmomanometer with the appropriate cuff size. The measurements for BP were taken by trained medical personnel at enrollment.

### Diet management for the GDM women

The initial treatment of GDM involves diet modification, glucose monitoring, and moderate exercise (15, 16). All the GDM participants in the study received 2 weeks of dietary management and nutritional recommendations at enrollment, which showed the guidelines for the subjects. Participants were considered as adhering to the given dietary recommendations in the presence of all the following criteria: carbohydrates 35–45% of total energy, rapidly absorbed sugars <10% of total energy, proteins 18–20% of total energy, fats 35% of total energy, fiber intake of at least 20–25 g/day, and no alcohol consumption. The nutritionist was in continuous contact with the enrolled GDM subjects, through weekly telephone contact, to remain updated regarding the nutritional condition of the subjects as the study progressed. Patients were instructed to self-monitor their blood glucose by finger-prick capillary blood glucose tests at least 4 times per day.

To reduce the effect of diet on the composition of the gut microbiota, general 2-week dietary restrictions were imposed on the healthy participants, including no peppery food and no yogurt intake and appropriate fat intake (the intake of calories from fat was no more than 35% of the total calories).

### Stool sample collection and DNA extraction

After providing written informed consent, all subjects were contacted for detailed instructions on how to collect and transport the stool sample. Stool samples of 57 subjects were collected at the time of enrollment for the first time. The second stool samples for GDM subjects were collected at the end of the study after the 2-week dietary intervention. For healthy pregnant women, the second stool samples were collected at the end of 2 weeks without dietary management intervention. Stool samples were self-collected by all the participants using the specimen collection kit as instructed. The fecal samples were collected at home, transferred to the hospital and immediately stored at −80 °C until DNA extraction. DNA was extracted from stool samples using the QIAamp DNA Stool Mini kit protocol (Qiagen, Germany). During the stool collection, one GDM sample at enrollment from one patient (G28) were limited, and the second sample was collected the other day, which changed the serial number to G28-2 at enrollment and G28-3 at the end of study.

### Illumina library generation

The V4 region of the 16S rRNA gene was amplified using 515F (5’-GTGCCAGCMGCCGCGGTAA −3’) and 806R (5’-GGACTACHVGGGTWTCTAAT −3’). The V4-specific primer regions were associated with the adaptor and the sequences, which were complementary to the Illumina forward and reverse sequencing primers. Each PCR product of the appropriate size was purified and quantified using a Qubit fluorometer and then added to a master pool of DNA for 250-bp nucleotide paired-end read assembly using the HiSeq 2500 genome analyzer (Illumina HiSeq 2500, USA).

### Bioinformatics

The RDP Classifier was used to assign all of the 16S rRNA gene sequences to a taxonomic hierarchy. The assembled reads were analyzed. The relative abundances of the various phyla, families and genera in each sample were computed and compared between the GDM patients and the healthy subjects. The trimmed reads were clustered into operational taxonomic units (OTUs) at 97 % identity. The comparison of the bacterial diversity of these samples was performed using the Chao1 richness index, ACE index and observed species. The reads displaying greater than 0.1% abundance in both groups were further analyzed via partial least-squares discriminant analysis (PLS-DA) to visualize the differences between two groups using the standard Simca-p1 software (version 12.0; http://www.umetrics.com/). The Principal Co-ordinates Analysis (PcoA) analyzed were performed based on Unweighted Unifrac distance metric.

### Statistical analysis

The microbial comparisons between the GDM and healthy groups were performed using the Mann-Whitney test. Associations between clinical indices and gut microbiota were evaluated by the Spearman rank correlation coefficient method. The difference in alpha-diversity between groups during GDM and non-GDM was assessed using Student’s t test. Statistical analysis of the clinical data was performed using SPSS (Statistical Package for Social Sciences) 22.0 software (SPSS Inc., Chicago, IL, USA). *P*<0.05 was considered significantly different.

### Availability of data

The raw sequences are available from the Genome Sequence Archive (GSA), the accession is: CRA004782.

## Results

### Characteristics of the patients

A flow chart illustrating the recruitment strategy of GDM and healthy subjects is shown in Fig 1. Clinical data from 27 GDM patients and 30 healthy controls are shown in Table 1. All 27 GDM patients and 30 healthy pregnant women were from the Peking University People’s Hospital. The mean age of the subjects was 32.7±3.3 years for the GDM group and 31.4 ±2.9 years for the healthy group. There were no differences in age or nulliparity rate between the two groups. The prepregnancy BMI value of the GDM group was 24.2±4.4, which was significantly higher than the value of 21.4±2.8 of the healthy group (*P=*0.0059), and the same trend was observed for the BMI at enrollment (27.1±4.3 vs. 25.0±2.9, GDM vs. healthy, *P=*0.038). The GDM group had a markedly higher systolic BP (SBP) value than that of the control group (mean 125.3±11.8 vs. 115.8±14.2, GDM vs. healthy, *P=*0.008), and an increased diastolic BP (DBP) value was found in GDM women compared to that of healthy women (mean 78.8±9.5 vs. 73.6±8.8, GDM vs. healthy, *P=*0.038). In the OGTT test, the GDM group had higher values at 0 h, 1 h and 2 h than the values of the healthy group (all *P*<0.001).

**Fig 1.**
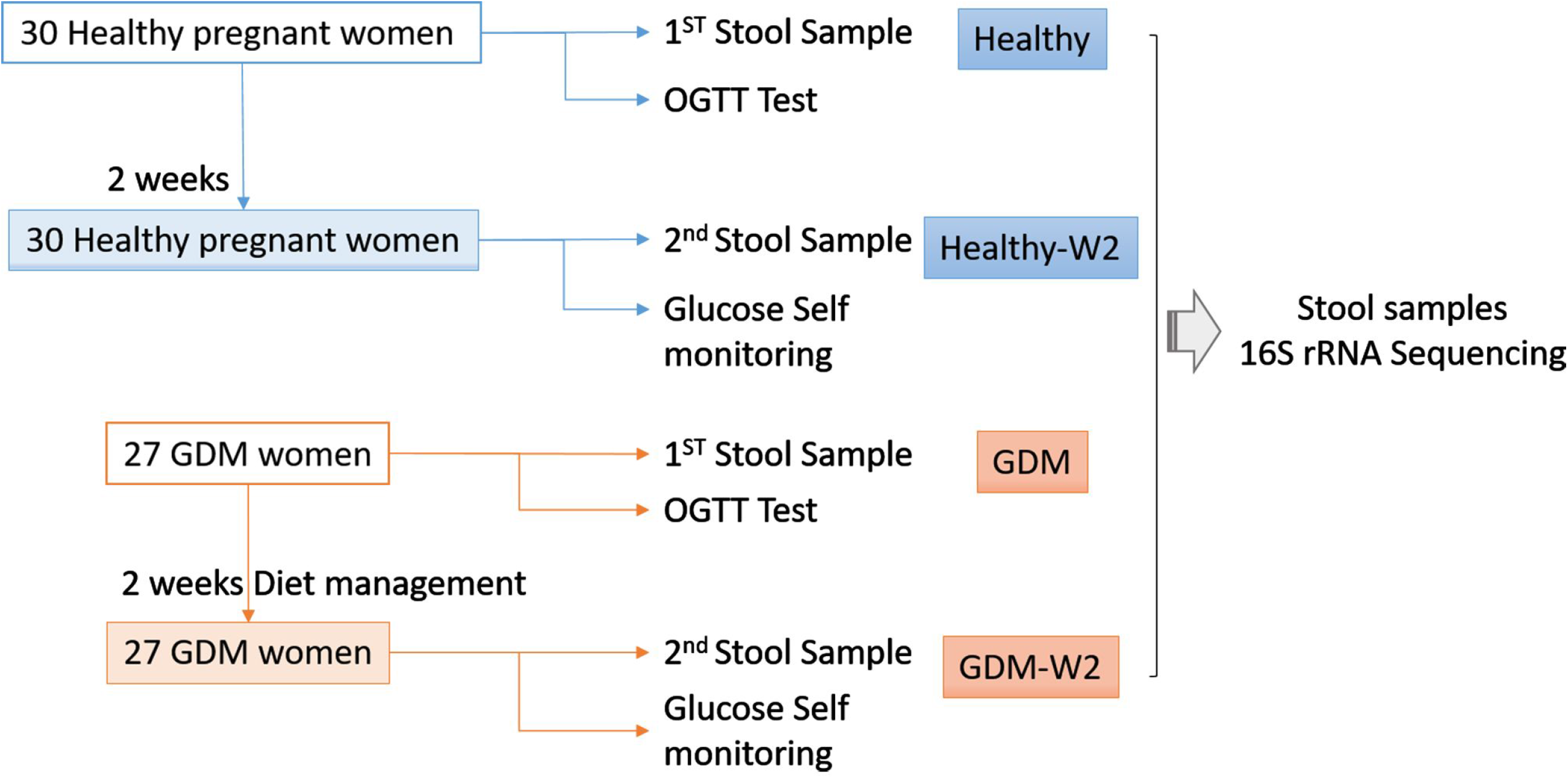
Flow chart illustrating the recruitment of GDM and healthy subjects.

**TABLE 1.** The clinical characteristics of all the GDM patients differ from those of the healthy participants

| | GDM<br>(Mean ± SD) | Healthy<br>(Mean ± SD) | *P* value |
| --- | --- | --- | --- |
| Number | 27 | 30 |  |
| Age | 32.7±3.3 | 31.4 ±2.9 | 0.11 |
| Prepregnancy weight (kg) | 63.5±12.2 | 57.3±8.9 | 0.031 |
| BMI (kg/m<sup>2</sup>) | 24.2±4.4 | 21.4±2.8 | 0.0059 |
| Enrollment weight (kg) | 71.1±12.4 | 66.9±9.5 | 0.15 |
| BMI (kg/m<sup>2</sup>) | 27.1±4.3 | 25.0±2.9 | 0.038 |
| Nulliparous (number) | 22/27 | 24/30 |  |
| Systolic BP (mmHg) | 125.3±11.8 | 115.8±14.2 | 0.008 |
| Diastolic BP (mmHg) | 78.8±9.5 | 73.6±8.8 | 0.038 |
| OGTT (mg/dL) |  |  |  |
| 0 min | 5.2±1.4 | 4.3±0.3 | 0.001 |
| 60 min | 10.1±1.6 | 7.3±1.4 | <0.0001 |
| 120 min | 8.8±1.3 | 6.4±1.2 | <0.0001 |

### Differences in fecal microbial communities between the healthy and GDM groups

To demonstrate the GDM microbiota signature, we explored the microbial composition of pregnant women with GDM. First, we performed PCoA using OTU relative abundance, and we observed discrete clustering of intestinal microbiota in the GDM and healthy groups at enrollment (Fig 2A). Additionally, shared or unique OTUs in the GDM and control groups were assessed to detect whether GDM has an effect on the gut microbiota. We found that the GDM group had more unique OTUs than the control group, with approximately 60.6% (1458/2404) unique OTUs compared with 14.3% (158/1104) in healthy women, signifying that GDM patients largely harbor unique inhabitant niches (Fig 2B).

**Fig 2.**
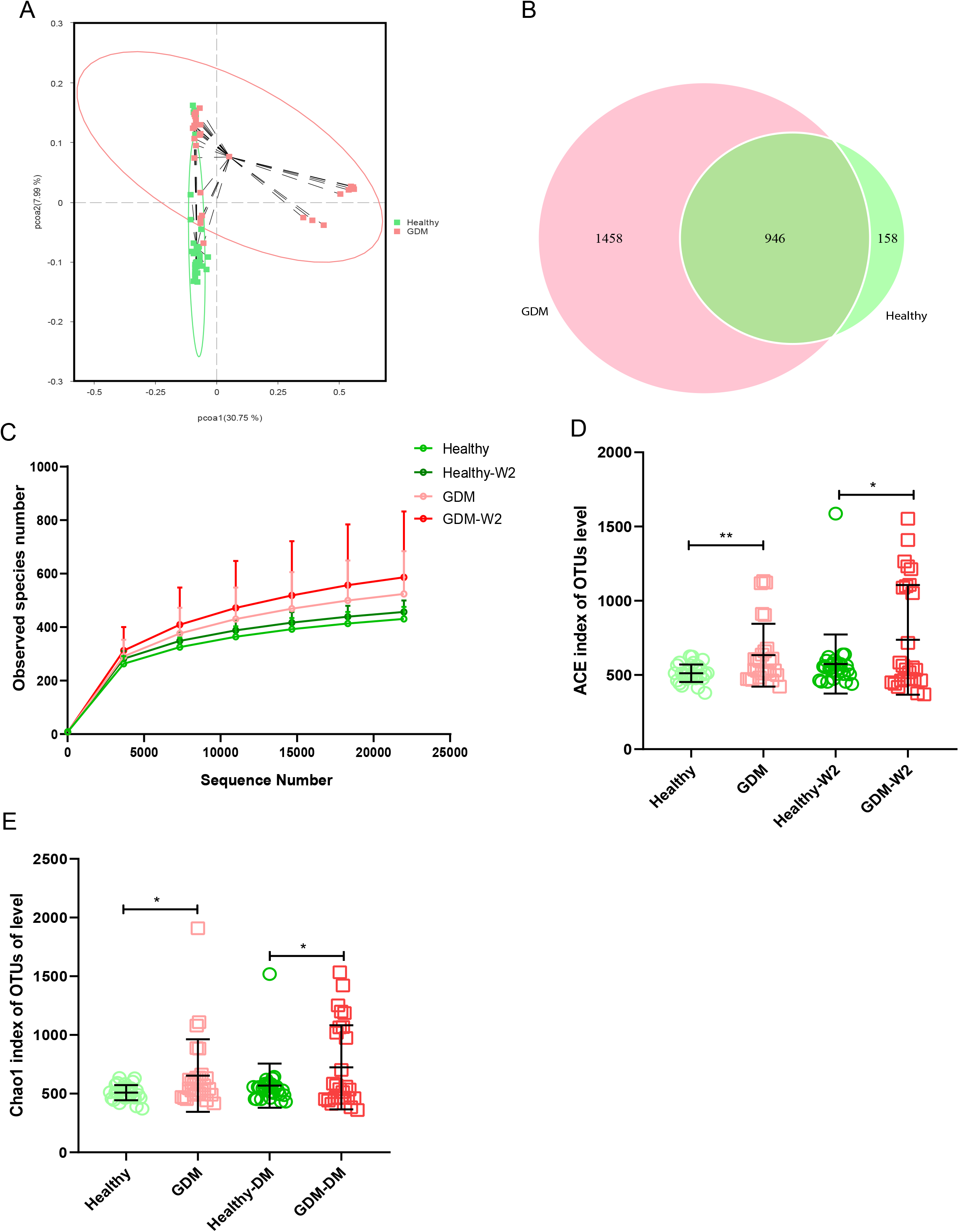
Comparison of the fecal microbiota composition between the GDM and healthy groups. **A.** Principal coordinate analysis (PCoA) at the OTU level between the GDM and healthy groups. **B.** Venn diagram illustrating the overlap of the OTUs identified in the fecal microbiota between the GDM and healthy groups. **C.** Observed species of 4 groups, including the GDM and healthy and the GDM-W2 and healthy-W2 groups. **D & E.** Alpha-diversity based on the ACE index and Chao 1 index at the OTU level. Mann-Whitney test, GDM vs. healthy, \*\**P*<0.01, \**P*<0.01.

The observed species of GDM samples were higher than non-GDM samples (Fig 2C). The ACE and Chao1 indices for alpha-diversity were both significantly increased in the GDM group (Fig 2D&2E), suggesting increased commensal diversity in GDM patients. Similar trends of alpha-diversity were also observed between the Healthy-W2 and GDM-W2 (diet management) groups, suggesting that the microbial pattern of women with GDM is distinct from that of healthy subjects at enrollment and at the end of the study.

### Microbiota structure of GDM patients based on taxonomic comparison

To further demonstrate these variations corresponding to the structure of the gut microbiota in GDM, we compared the bacterial abundance between groups at the phylum level (Fig 3A). No significant differences were observed between the healthy subjects and the GDM subjects at enrollment for most of the phyla, with the exception of *Acidobacteria*, which was found to be 0.51% in the GDM group compared with 0.37% in the healthy group (*P*=0.001). The microbial compositions at the phylum level for each sample at enrollment and at the end of the study are shown in Fig S1. Interestingly, *Acidobacteria* was associated with increased levels of blood glucose in the 0-h OGTT (Fig 3B).

**Fig 3.**
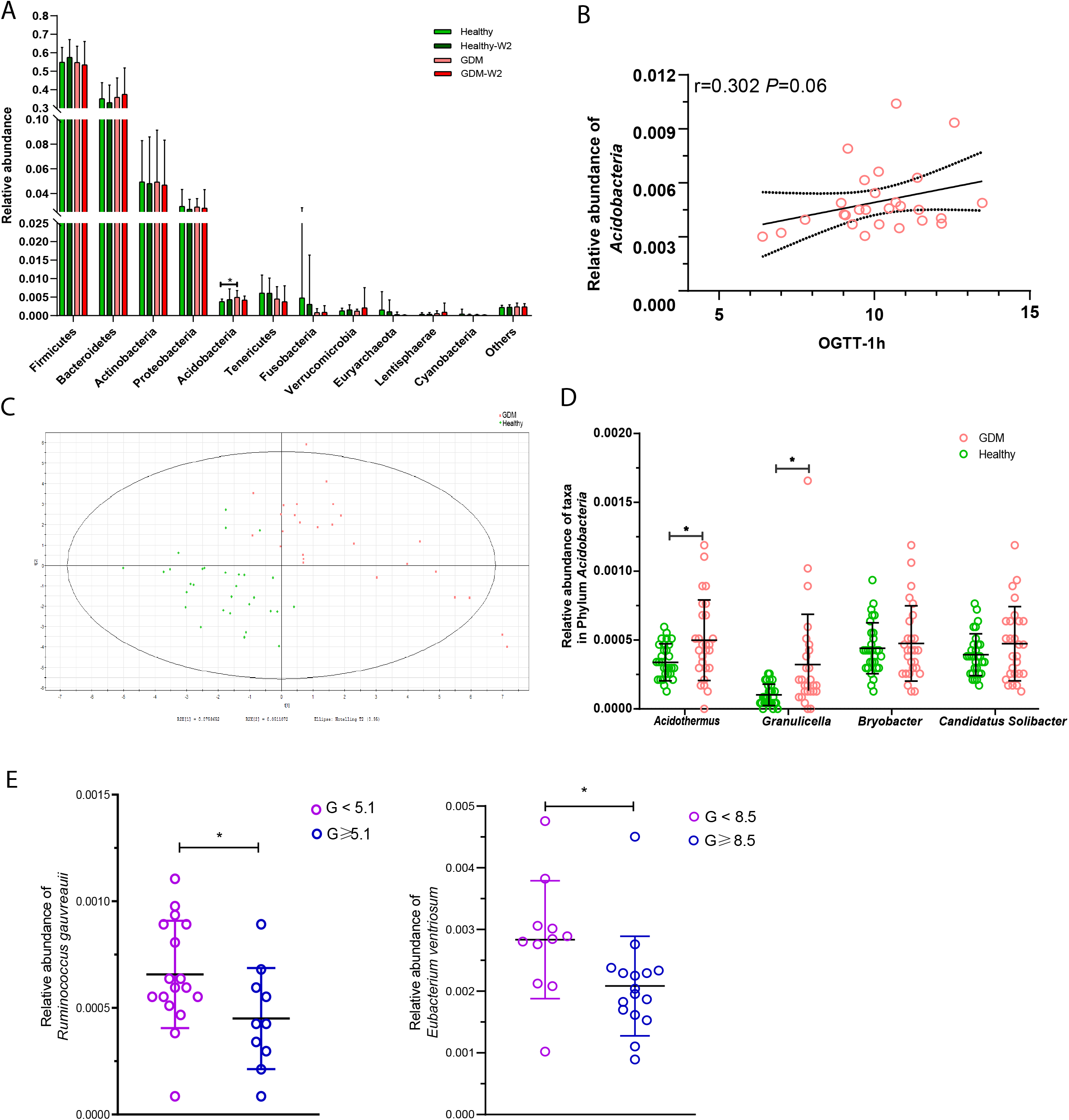
Abundances of taxa in GDM and healthy participants. **A.** Comparison of the relative abundances at the phylum level among the four GDM and non-GDM groups. The Mann–Whitney test was used to evaluate the two groups. \**P*<0.05. **B.** PLS-DA score plots based on the relative abundances of microbiota between the GDM and healthy groups. **C.** Correlation between the relative abundance of the phylum *Acidobacteria* and the 1-h OGTT measurement. Spearman analysis, R=0.302, *P*=0.06. **D.** Comparison of the relative abundances of *Acidothermus, Granulicella, Bryobacter*, and *Candidatus_Solibacter* in the phylum *Acidobacteria* in the GDM and healthy groups. Mann-Whitney test, GDM vs. control, \*\**P*<0.01, \**P*<0.01. **E.** The relative abundances of *Ruminococcus gauvreauii* and *Eubacterium ventriosum* were highly correlated with the OGTT values at 0 h and 2 h. Mann-Whitney test, GDM vs. healthy, \*\**P*<0.01, \**P*<0.01.

Next, we compared taxa at the genus level. The PLS-DA method was performed (Fig 3C). Forty-nine key genera with variable importance in projection (VIP) scores >1 were identified that differentiated the GDM and healthy groups (Table 2). We then clustered the samples according to the relative abundance of the 49 genera. Twenty-seven genera were enriched in the GDM microbiota samples, with 4 genera (*Acidothermus, Granulicella, Bryobacter*, and *Candidatus_Solibacter*) belonging to the phylum *Acidobacteria*. Among them, *Acidothermus* and *Granulicella* were significantly enriched in the GDM group (Fig 3D). Seven genera belonging to *Proteobacteria*, including *Citrobacter, Burkholderia, Acidibacter*, and *Bilophila*, were significantly highly expressed in the GDM intestinal microbiota (*P*<0.05). The genera *Eubacterium*, *Holdemania*, and *Tyzzerella*, in the phylum *Firmicutes*, were rarely detected in women with healthy pregnancy microbiota compared with women with GDM. The remaining 22 genera of the 49 key phylotypes were overexpressed in healthy pregnant microbiota, some of which even disappeared in GDM patients. One genus, *Ruminococcaceae_UCG-010*, belonging to *Firmicutes*, was highly enriched in the healthy group. Additionally, *Akkermansia* (*P*=0.067) and *Coprococcus_2* (*P*=0.027) were increased in healthy subjects. *Akkermansia* was recently proven to be a crucial player in maintaining the integrity of the gastrointestinal tract. In nonpregnant adults with metabolic syndrome and type 2 diabetes, *Akkermansia* is reported to be depleted as well (17–19). Our findings suggest that the gut microbiota of women with GDM has similarities with the microbiota reported in patients with type 2 diabetes and associated intermediary metabolic traits. At the OTU level, a reduced abundance of *Akkermansia* has previously been reported in the third trimester of healthy pregnant women (20).

**TABLE 2.** Forty-nine key genera with VIP >1 that were differentially expressed in the GDM and healthy groups

| Genus with VIP $\geq 1$ | GDM<br>mean | Healthy<br>mean | GDM/Healt<br>hy | *P*<br>value | Phylum |
| --- | --- | --- | --- | --- | --- |
| *Citrobacter* | 0.000316 | 2.41E-05 | up | 0.048 | *Proteobacteria* |
| *Bradyrhizobium* | 0.000422 | 8.64E-05 | up | 0.065 | *Proteobacteria* |
| *Eubacterium* | 0.00012 | 3.4E-05 | up | 0.001 | *Firmicutes* |
| *Granulicella* | 0.000323 | 0.000102 | up | 0.001 | *Acidobacteria* |
| *Holdemania* | 0.000187 | 7.5E-05 | up | 0.014 | *Firmicutes* |
| *Succinivibrio* | 8.81E-05 | 3.54E-05 | up | 0.212 | *Proteobacteria* |
| *Oscillibacter* | 0.000211 | 9.06E-05 | up | 0.44 | *Firmicutes* |
| *Tyzzerella* | 0.000856 | 0.000368 | up | 0.007 | *Firmicutes* |
| *Holdemanella* | 0.009481 | 0.004176 | up | 0.162 | *Firmicutes* |
| *Paraprevotella* | 0.000994 | 0.000578 | up | 0.126 | *Bacteroidetes* |
| *Victivallis* | 0.00056 | 0.000344 | up | 0.042 | *Lentisphaerae* |
| *Desulfovibrio* | 0.000458 | 0.00029 | up | 0.479 | *Proteobacteria* |
| *Lachnospiraceae* | 0.002291 | 0.001517 | up | 0.137 | *Firmicutes* |
| *Burkholderia* | 0.000824 | 0.000551 | up | 0.027 | *Proteobacteria* |
| *Acidothermus* | 0.000499 | 0.000338 | up | 0.034 | *Acidobacteria* |
| *Acidibacter* | 0.000677 | 0.000508 | up | 0.405 | *Proteobacteria* |
| *Mucilaginibacter* | 0.00037 | 0.00028 | up | 0.02 | *Bacteroidetes* |
| *Candidatus_Solibacter* | 0.000474 | 0.000394 | up | 0.404 | *Acidobacteria* |
| *Ruminiclostridium_9* | 0.001163 | 0.00098 | up | 0.141 | *Firmicutes* |
| *Ruminococcus_gauvreauii* | 0.000581 | 0.000491 | up | 0.214 | *Firmicutes* |
| *unidentified_Ruminococcacea* |  |  |  |  |  |
| *e* | 0.001548 | 0.001359 | up | 0.949 | *Firmicutes* |
| *Roseburia* | 0.028429 | 0.025656 | up | 0.482 | *Firmicutes* |
| *Bilophila* | 0.002439 | 0.002216 | up | 0.179 | *Proteobacteria* |
| *Alistipes* | 0.011983 | 0.010959 | up | 0.354 | *Bacteroidetes* |
| *Bryobacter* | 0.000475 | 0.00044 | up | 0.968 | *Acidobacteria* |
| *Odoribacter* | 0.001495 | 0.001395 | up | 0.302 | *Bacteroidetes* |
| *Dorea* | 0.007676 | 0.007233 | up | 0.678 | *Firmicutes* |
| *Lachnospiraceae_NK4A136* | 0.002712 | 0.002715 | down | 0.438 | *Firmicutes* |
| *Eubacterium_ruminantium* | 0.00633 | 0.006883 | down | 0.26 | *Firmicutes* |
| *Bifidobacterium* | 0.033865 | 0.038103 | down | 0.56 | *Acidobacteria* |
| *Ruminococcaceae_UCG-013* | 0.001251 | 0.00142 | down | 0.994 | *Firmicutes* |
| *Tyzzerella_3* | 0.002118 | 0.002482 | down | 0.073 | *Firmicutes* |
| *Ruminococcaceae_UCG-005* | 0.00281 | 0.003448 | down | 0.452 | *Firmicutes* |
| *Ruminococcaceae_UCG-002* | 0.005792 | 0.00715 | down | 0.056 | *Firmicutes* |
| *Ruminococcaceae_NK4A214* | 0.00144 | 0.001797 | down | 0.083 | *Firmicutes* |
| *Eubacterium_ventriosum* | 0.00239 | 0.003046 | down | 0.207 | *Firmicutes* |
| *Enterococcus* | 0.001193 | 0.001627 | down | 0.09 | *Firmicutes* |
| *Megasphaera* | 0.001971 | 0.00306 | down | 0.749 | *Firmicutes* |
| *Lachnospiraceae_UCG-003* | 0.000269 | 0.000419 | down | 0.11 | *Firmicutes* |
| *Coprococcus_2* | 0.005271 | 0.008583 | down | 0.027 | *Firmicutes* |
| *Ruminiclostridium_5* | 0.001904 | 0.003111 | down | 0.009 | *Firmicutes* |
| *Ruminococcaceae_UCG-010* | 0.000848 | 0.001403 | down | 0 | *Firmicutes* |
| *Sarcina* | 0.000126 | 0.000252 | down | 0.001 | *Firmicutes* |
| *Butyrivibrio* | 0.000455 | 0.000927 | down | 0.02 | *Firmicutes* |
| *Intestinimonas* | 3.93E-05 | 8.64E-05 | down | 0.07 | *Firmicutes* |
| *Akkermansia* | 0.000189 | 0.000435 | down | 0.067 | *Verrucomicrobia* |
| *Weissella* | 7.87E-05 | 0.000217 | down | 0.002 | *Firmicutes* |
| *Prevotella_2* | 0.001153 | 0.003598 | down | 0.108 | *Bacteroidetes* |

To further examine the relationship between these VIP genera in GDM, we evaluated their abundance based on the results of the OGTT. The threshold values (5.1 at 0 h, 10.0 at 1 h and 8.5 at 2 h during the OGTT) are based on the diagnostic criteria recommended by the International Association of the Diabetes and Pregnancy Study Groups in 2011. As shown in Fig 3E, two short chain fatty acids producing and anti-inflammatory bacteria were highly correlated with the OGTT value at 0 h and 2 h. The relative abundance of *Ruminococcus gauvreauii* was observed depleted in GDM women with abnormal OGTT value at 0 h (*P=*0.046), and the relative abundance of *Eubacterium ventriosum* was decreased in GDM women with the abnormal OGTT value at 2 h (*P=*0.009, Mann-Whitney test).

### Microbiota signature after dietary intervention

We found that GDM patients developed a microbial pattern with higher alpha-diversity after diet management (Fig 2D & E). Compared with the GDM samples, the GDM-W2 samples showed some distinct taxa with VIP scores >1, according to the PLS-DA analysis (Fig S2).

At the family level, GDM-W2 samples showed decreased pathogenic taxa (*Acidaminococcaceae*, *Enterobacteriaceae*, and *Bacteroidaceae*) and increased *Bifidobacteriaceae* and butyric acid-producing bacteria (*Prevotellaceae* and *Lachnospiraceae*) compared with the GDM microbial samples at enrollment, suggesting a better pattern driven by the 2 weeks of diet management. One more interesting observation is that because the bacterial lineages were constant within pregnancy over time, communities from the same GDM person were generally more similar to one another than to those from other people from the healthy group (Fig 4B).

**Fig 4.**
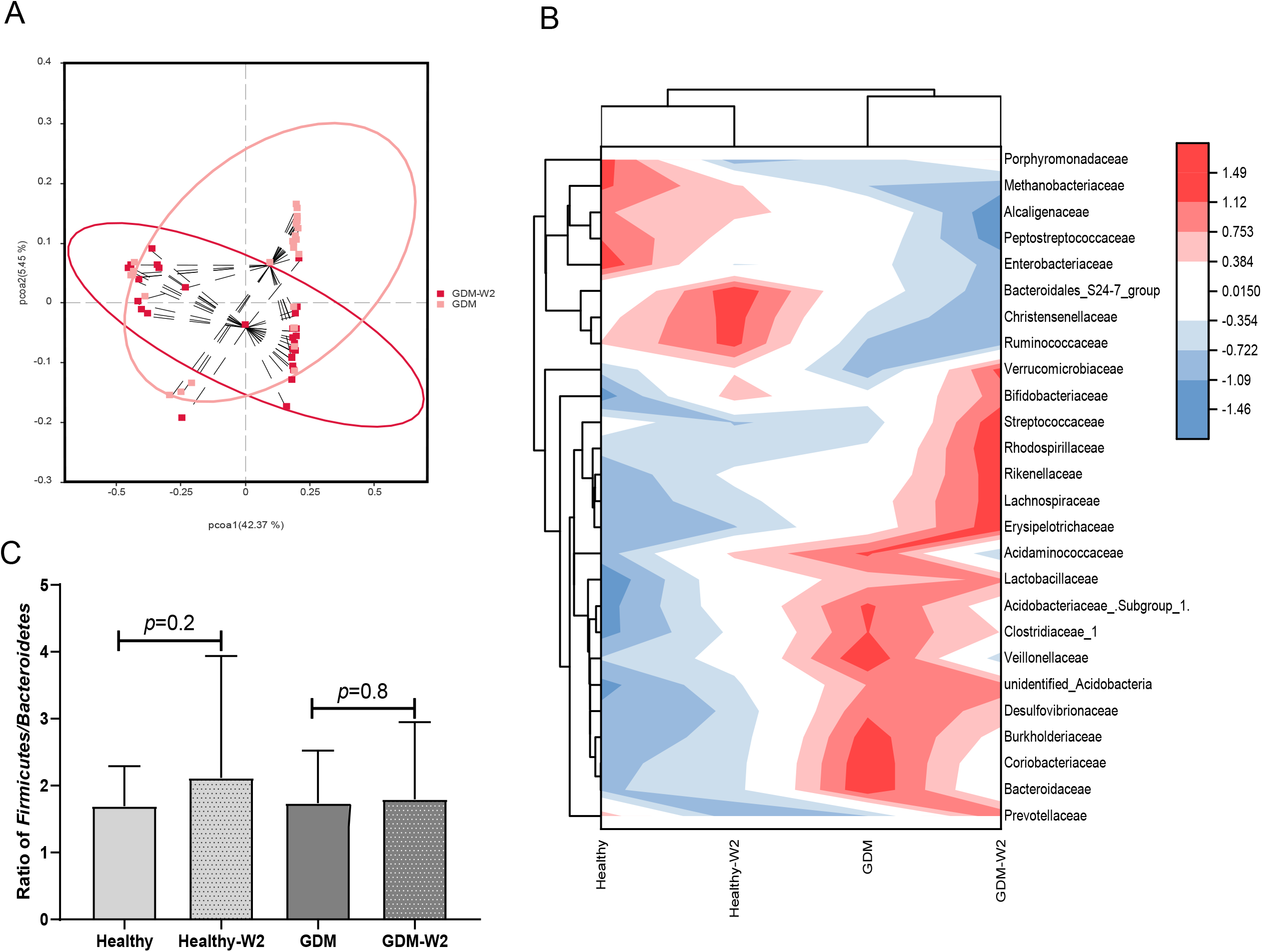
The microbial pattern after diet management. **A.** Principal coordinate analysis (PCoA) at the OTU level between the GDM-W2 and healthy-W2 groups. **B.** Heatmap analysis of the differentially expressed taxa at the family level. **C.** Ratio of *Firmicutes/Bacteroidetes* among the GAM and non-GDM groups with or without diet intervention.

It is presumed that the influence of maternal gestational diet on the phylogenetic structure of the intestinal microbiota during pregnancy remains underexplored in well-controlled models. To investigate whether the microbiota can be driven by dietary management for GDM in pregnancy, the two dominant groups of beneficial bacteria, *Bacteroidetes* and *Firmicutes*, were analyzed. At the phylum level, a slightly increase in the *Firmicutes*/*Bacteroidetes* (F/B) ratio in late pregnancy was exhibited in the GDM group compared with the non-GDM group (Fig 4C). Previous studies indicated that a higher *Firmicutes*/*Bacteroidetes* ratio was associated with obesity (21) and an aggravation of low-grade inflammation (22). Here, we showed that after 2 weeks of diet therapy, the relative abundance of *Bacteroidetes* in GDM samples increased, and the abundance of *Firmicutes* decreased slightly (Fig 2A). More importantly, the ratio of *Firmicutes*/*Bacteroidetes* did not increase in GDM-W2 fecal samples compared with GDM samples at enrollment (*P*=0.8) (Fig 4C). However, without diet management, an obviously increased proportion of *Firmicutes/Bacteroidetes* (*P=*0.2) developed in healthy pregnancies (healthy-W2 samples).

Four genera (*Acidothermus, Granulicella, Bryobacter*, and *Candidatus_Solibacter*) belonging to the phylum *Acidobacteria* were increased in the GDM group, compared with healthy group. Furthermore, we evaluated the levels of the 4 genera in GDM with dietary management (Fig S3). A total of 66.7% (18/27) of GDM subjects showed decreased levels of the genus *Acidothermus* after 2 weeks of diet management. In contrast, 59.3% (16/27) of GDM samples showed decreased levels of the genera *Granulicella, Bryobacter*, and *Candidatus Solibacter* after 2 weeks of diet management.

### Association between fecal microbiota and clinical parameters

We examined the correlations between the OGTT values (0 h, 1 h and 2 h), BMI indices (prepregnancy and at enrollment), blood pressure values (SBP and DBP) and the genera of the fecal microbiota (Fig 5).

**Fig 5.**
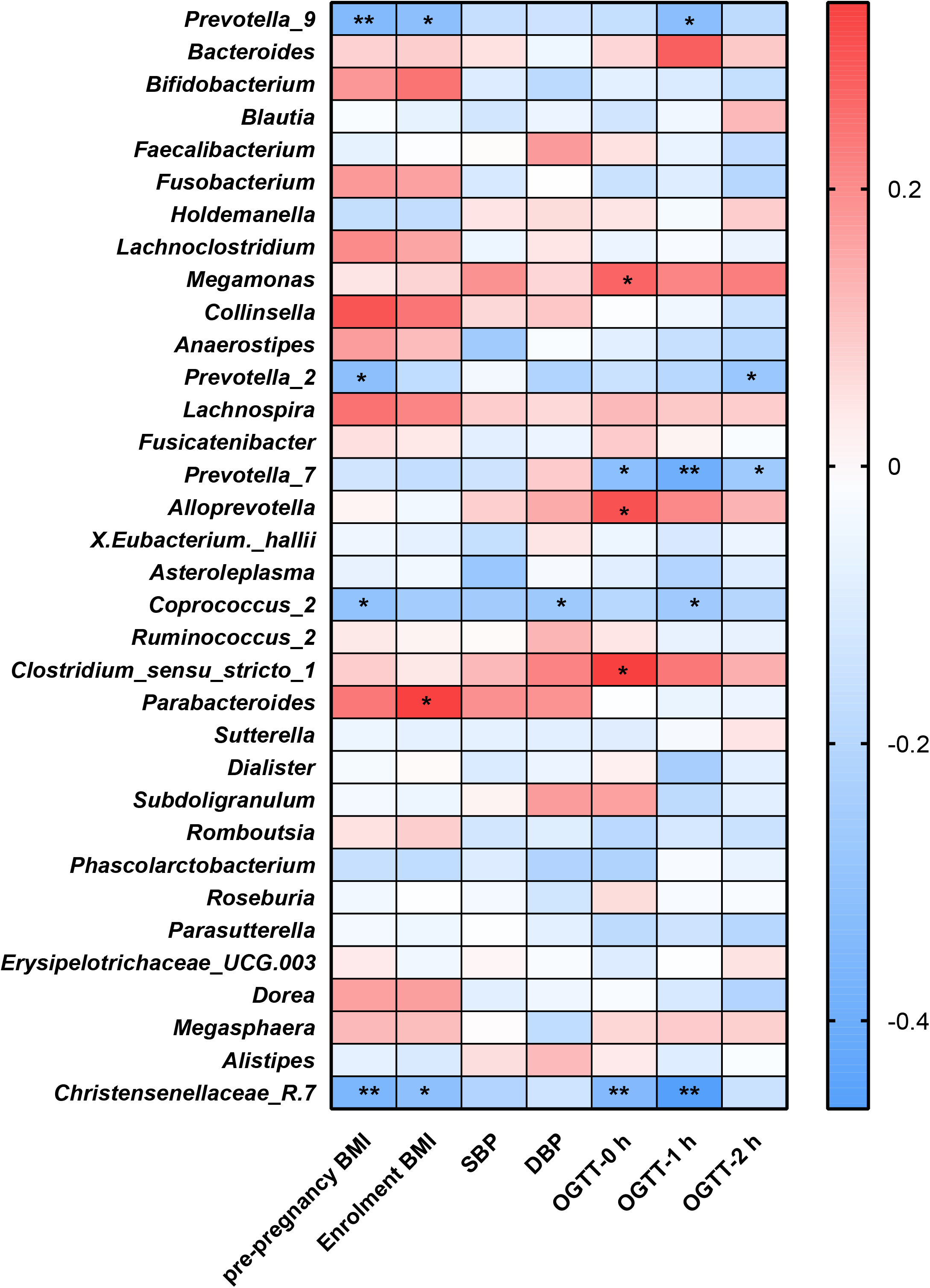
Heatmap analysis of the correlation between the gut microbiota composition and clinical scores.

The distribution of correlation coefficients by heatmap analysis showed that the *Coprococcus_2, Christensenellaceae*_R.7, and *Prevotella* groups (*Prevotella_2, Prevotella_7* and *Prevotella_9*) were negatively correlated with the OGTT value, BP values and BMI index (*P*<0.05); among them, *Coprococcus_2* was significantly increased in the healthy group compared with the GDM group.

*Parabacteroides* showed positive correlations with BMI at enrollment (*P*<0.05). Additionally, *Alloprevotella, Megamonas* and *Clostridium_sensu_stricto-1* showed positive correlations with GDM-correlated clinical measures and OGTT values at 0 h (*P*<0.05). Previous studies observed that the genus *Megamonas* was increased in GDM patients in late pregnancy. Elevated genera of *Megamonas* have also been reported to be associated with higher blood glucose at an individual level (9, 23–25).

## Discussion

Studies support a causal role for the gut microbiota in the development of type 2 diabetes, insulin resistance and obesity (26). In this study, we compared the composition of the human intestinal microbiota between GDM patients and healthy subjects using a culture-independent Illumina HiSeq 2500 platform. The aim of the present study was to identify gut microbiota dysbiosis in GDM subjects and the associated microbial changes in GDM-W2 samples after diet intervention for 2 weeks and compare them with the basal GDM microbial composition. We observed a marked shift in the microbiota composition at the phylum and genus levels in GDM samples compared with healthy samples and identified the microbial pattern of GDM-W2 samples after a 2-week dietary intervention.

Gut dysbiosis in women with GDM was mainly characterized by changes in microbiota diversity. It was previously reported that an increase was found in the alpha-diversity in the third trimester of GDM women when compared to the level of the control group (24). Regarding alpha-diversity, we used the ACE and Chao1 indices and found significant separation in the alpha-diversity between GDM and non-GDM individuals at their enrollment and at the end of the study, indicating dysbiosis of the gut microbiota in GDM women compared with healthy pregnant women. To further identify gut microbial dynamics, the different bacterial taxa were compared within the GDM and non-GDM groups. At the phylum level, the abundance of *Acidobacteria* was significantly greater in the gut microbiota of GDM samples and was associated with increased levels of blood glucose in the 0-h OGTT (Fig 3B). In particular, we observed significant elevation of *Acidothermus* and *Granulicella* belonging to the phylum *Acidobacteria* in the GDM group. The phylum *Acidobacteria* was reported in the gut microbiome of obese individuals (27) and was shown to contain a host of genes involved in diverse metabolic pathways, as evidenced by their pan-genomic profiles in the soil microbiota (28). Further exploration of these genetic attributes and more in-depth insights into GDM mechanics and dynamics would lead to a better understanding of the functions and biological significance of this elevated phylum in the GDM gut environment.

Several bacterial groups at the genus level were detected to be different in the GDM and healthy groups, such as *Megamonas* assigned to the phylum *Firmicutes*. The relationships between gastrointestinal *Megamonas* and metabolic disorders such as obesity and type 2 diabetes have recently been discovered (29). Differential abundance testing showed that *Megamonas, Bacteroides*, and *Eubacterium* were statistically associated with food addition (30). A recent study also suggested that the abundance of *Megamonas*, which is closely related to childhood obesity, increased in the gut microbiota of obese children (29). Of particular interest, we revealed the association between gut *Megamonas* and GDM. Our results showed that *Megamonas* was positively correlated with higher blood glucose in the OGTT test at 0 h in the GDM samples at enrollment (Fig 5). Members of *Megamonas* are known to produce acetic and propionic acid, which is beneficial for the balance of glucose uptake (31). Systemic disorders of glucose metabolism might be modulated by the related gut microbiota. Further study to explore the composition of *Megamonas* and the production of metabolites involved in glucose homeostasis *in vitro* and *in vivo* is very important.

Short-chain fatty acids (SCFAs), especially acetate, propionate and butyrate, are the end products of the intestinal microbial fermentation of dietary fibers and resistant starch. It is well documented that plasma and colonic SCFAs are associated with metabolic syndromes, i.e., obesity and type 2 diabetes (32). SCFAs, namely, acetate, butyrate, and propionate, have been reported to affect metabolic activities at the molecular level. Acetate affects the metabolic pathway through the G protein-coupled receptor (GPCR) and free fatty acid receptor 2 (FFAR2/GPR43). The FFAR2 signaling pathway regulates insulin-stimulated lipid accumulation in adipocytes and inflammation (33, 34). *Coprococcus_2*, an acetate-producing bacteria (25, 35), was found to be negatively correlated with the OGTT value at 1 h, BP values and prepregnancy BMI index (*P*<0.05) by Spearman analysis and was significantly higher in the healthy group than in the GDM group. *Coprococcus* was also proven to be altered in the fecal microbiota of patients with polycystic ovary syndrome, which is a metabolic disorder (36). Guo et al. (37) found that *Coprococcus* deletion is implicated in many of the outcomes, including glucose homeostasis. The importance of an association between the deletion of the *Coprococcus* genus and high levels of blood glucose at 1-h in the OGTT measure is therefore supported by the acetate-producing effect. Furthermore, other SCFA-producing taxa, including *Prevotella_2, Prevotella_7*, and *Prevotella_9*, were found to be negatively associated with OGTT measures and the BMI index separately, indicating a beneficial effect on blood glucose in GDM subjects (38). We presumed that acetate arising from *Coprococcus_2* members and succinate from *Prevotalla* members are important for energy metabolism and have a mainly protective role in relation to healthy pregnancy. Thus, the observed absence of the *Coprococcus_2* and *Prevotella* groups in the fecal microbiota of GDM could be a possible microbial driving force for GDM. A better understanding of the microbial ecology of colonic acetate- and succinate-producing bacteria, especially the *Coprococcus_2* and *Prevotella* groups, may help to explain the influence of diet on the acetate and succinate supply and may contribute to the development of new approaches for optimizing microbial activity for diet management for GDM subjects. *Eubacterium ventriosum*, another SCFAs producer, had been found negative correlated with visceral fat area (VFA) (39). Moraes et al. reported that the abundance of *E. ventriosum* were associated to better cardiometabolic profile (40). Consistent with our study, the data demonstrated a significant decrease of gut *Eubacterium ventriosum* from GDM subjects with abnormal OGTT values at 2 h (Fig 3E). Combined with these findings, we presumed that the expression of the SCFAs producers are critical for energy homeostasis during pregnancy. Further studies investigating the targets and signaling pathways of SCFAs in the GDM microbial, and the modulation of SCFAs-producing bacteria by diet intervention would benefit for GDM management.

Therefore, to further identify the role of diet intervention during GDM pregnancy, we analyzed the ratio of *Firmicutes/Bacteroidetes*, and a higher ratio was proposed as an eventual biomarker of obesity and other metabolic syndromes compared with normal-weight individuals (41). Our data showed different increases in the *Firmicutes/Bacteroidetes* ratio between the GDM and non-GDM groups. Healthy W2 samples without diet management showed a nearly significant increase in the *Firmicutes/Bacteroidetes* ratio, indicating a change in energy homeostasis during pregnancy. Similar to our findings on the *Firmicutes/Bacteroidetes* ratio in healthy pregnant women, Zheng et al. (42) reported that there were elevations in the *Firmicutes/Bacteroidetes* ratio in the second (T2) trimester compared with the first (T1) trimester. Ley et al. (22) reported that the *Firmicutes/Bacteroidetes* ratio decreases with weight loss on a low-calorie diet. In our observations, the *Firmicutes/Bacteroidetes* ratio did not change in GDM-W2 samples under diet management compared to the ratio in GDM samples, suggesting that the diet intervention could play a positive role during GDM pregnancy by affecting *Firmicutes/Bacteroidetes* ratio. In particular, the gut microbial pattern was not altered in the GDM group with or without 2 weeks of diet intervention (Fig 4A&B). In agreement with our observation, a controlled-feeding study showed that enterotype identity remained stable during the 10-day study, and alternative microbial states were associated with a long-term diet (43). Thus, we presume that the role of short-term diet management in GDM processes is associated with changes in the *Firmicutes/Bacteroidetes* ratio and some specific taxa rather than an alternative gut microbial pattern.

It is well suggested that the diet contributes to the gut microbiota composition in GDM (42). Microbiota-derived metabolites affect glucose homeostasis through intestinal gluconeogenesis (38). A few studies have examined the gut microbiota of GDM and healthy pregnant women before and after diet invention. Uniquely, in the present study, we could compare gut microbiota in GDM fecal samples, allowing identification of taxa that exhibited differential abundance at the two time points. We discovered that a short-term diet had a beneficial effect on GDM by modulating the *Firmicutes/Bacteroidetes* ratio and some taxa. This first observation of the high expression of the phylum *Acidobacteria* in GDM offered an important clue for further study on the subgroup of *Acidobacteria* and the mechanism of GDM. Several limitations in our study should be considered. One was that we did not have fecal samples after long-term dietary management. Additionally, our suggestion of the occurrence of specific taxa with divergent metabolites calls for future metagenomic sequencing to reveal the metabolic pathways of the key taxa. In conclusion, our results highlight the relevance of characterizing gut microbial population differences and contribute to understanding the plausible link between diet and specific gut bacterial species that are able to influence metabolic homeostasis and GDM development. Modulating the gut microbiota via short-term diet intervention, especially SCFA-producing bacteria, could be a promising strategy in the search for alternatives for the treatment of metabolic disorders in GDM (44–46). Long-term observation may be more valuable to study the dynamic alteration of the GDM gut microbiota.

## Acknowledgments

We thank all the subjects who made this study possible.

---

**Fig S1.** Comparison of the relative abundance at the phylum level between the 27 GDM and 30 healthy individuals at the time of enrolment and study end.

**Fig S2.** PLS-DA analysis indicated 49 distinct taxa with VIP score>1 between GDM samples and GDM-W2 samples. Mann-Whitney test, GDM vs. Healthy, \*\**P*<0.01, * *P*<0.01.

**Fig S3.** The *Acidothermus, Granulicella, Bryobacter, Candidatus_Solibacter* belonging to the phylum *Acidobacteria* were evaluated in GDM and GDM-W2 samples. The 66.7% (18/27) GDM samples was showed decreased level of genus *Acidothermus* after two-week diet management. While 59.3% (16/27) GDM samples was showed decreased level of genus *Granulicella, Bryobacter, Candidatus_Solibacter* after two-week diet management.

## References

1. Savage DC. 1977. Microbial ecology of the gastrointestinal tract. Annu Rev Microbiol 31:107–33.

2. Newbern D, Freemark M. 2011. Placental hormones and the control of maternal metabolism and fetal growth. Curr Opin Endocrinol Diabetes Obes 18:409–16.

3. Koren O, Goodrich JK, Cullender TC, Spor A, Laitinen K, Backhed HK, Gonzalez A, Werner JJ, Angenent LT, Knight R, Backhed F, Isolauri E, Salminen S, Ley RE. 2012. Host remodeling of the gut microbiome and metabolic changes during pregnancy. Cell 150:470–80.

4. Cani PD, Delzenne NM. 2007. Gut microflora as a target for energy and metabolic homeostasis. Curr Opin Clin Nutr Metab Care 10:729–34.

5. Vijay-Kumar M, Aitken JD, Carvalho FA, Cullender TC, Mwangi S, Srinivasan S, Sitaraman SV, Knight R, Ley RE, Gewirtz AT. 2010. Metabolic syndrome and altered gut microbiota in mice lacking Toll-like receptor 5. Science 328:228–31.

6. Juan J, Yang H. 2020. Prevalence, prevention, and lifestyle intervention of gestational diabetes mellitus in China. Int J Environ Res Public Health 17:9517.

7. Scheithauer TPM, Rampanelli E, Nieuwdorp M, Vallance BA, Verchere CB, van Raalte DH, Herrema H. 2020. Gut Microbiota as a Trigger for Metabolic Inflammation in Obesity and Type 2 Diabetes. Front Immunol 11:571731.

8. Turnbaugh PJ, Ley RE, Mahowald MA, Magrini V, Mardis ER, Gordon JI. 2006. An obesity-associated gut microbiome with increased capacity for energy harvest. Nature 444:1027–31.

9. Crusell MKW, Hansen TH, Nielsen T, Allin KH, Ruhlemann MC, Damm P, Vestergaard H, Rorbye C, Jorgensen NR, Christiansen OB, Heinsen FA, Franke A, Hansen T, Lauenborg J, Pedersen O. 2018. Gestational diabetes is associated with change in the gut microbiota composition in third trimester of pregnancy and postpartum. Microbiome 6:89.

10. Wang J, Zheng J, Shi W, Du N, Xu X, Zhang Y, Ji P, Zhang F, Jia Z, Wang Y, Zheng Z, Zhang H, Zhao F. 2018. Dysbiosis of maternal and neonatal microbiota associated with gestational diabetes mellitus. Gut 67:1614–1625.

11. Buchanan TA, Xiang AH, Page KA. 2012. Gestational diabetes mellitus: risks and management during and after pregnancy. Nat Rev Endocrinol 8:639–49.

12. Johnson AJ, Vangay P, Al-Ghalith GA, Hillmann BM, Ward TL, Shields-Cutler RR, Kim AD, Shmagel AK, Syed AN, Personalized Microbiome Class Students, Walter J, Menon R, Koecher K, Knights D. 2019. Daily sampling reveals personalized diet-microbiome associations in humans. Cell Host Microbe 25:789–802.e5.

13. Bassis CM. 2019. Live and Diet by Your Gut Microbiota. mBio 10.

14. Ferrocino I, Ponzo V, Gambino R, Zarovska A, Leone F, Monzeglio C, Goitre I, Rosato R, Romano A, Grassi G, Broglio F, Cassader M, Cocolin L, Bo S. 2018. Changes in the gut microbiota composition during pregnancy in patients with gestational diabetes mellitus (GDM). Sci Rep 8:12216.

15. Blumer I, Hadar E, Hadden DR, Jovanovic L, Mestman JH, Murad MH, Yogev Y. 2013. Diabetes and pregnancy: an endocrine society clinical practice guideline. J Clin Endocrinol Metab 98:4227–49.

16. American Diabetes Association. 2014. Standards of medical care in diabetes--2014. Diabetes Care 37:S14–S80.

17. Hills RD, Jr., Pontefract BA, Mishcon HR, Black CA, Sutton SC, Theberge CR. 2019. Gut Microbiome: Profound Implications for Diet and Disease. Nutrients 11.

18. Everard A, Belzer C, Geurts L, Ouwerkerk JP, Druart C, Bindels LB, Guiot Y, Derrien M, Muccioli GG, Delzenne NM, de Vos WM, Cani PD. 2013. Cross-talk between Akkermansia muciniphila and intestinal epithelium controls diet-induced obesity. Proc Natl Acad Sci U S A 110:9066–71.

19. Macchione IG, Lopetuso LR, Ianiro G, Napoli M, Gibiino G, Rizzatti G, Petito V, Gasbarrini A, Scaldaferri F. 2019. Akkermansia muciniphila: key player in metabolic and gastrointestinal disorders. Eur Rev Med Pharmacol Sci 23:8075–8083.

20. Yao Z, Long Y, Ye J, Li P, Jiang Y, Chen Y. 2020. 16S rRNA Gene-Based Analysis Reveals the Effects of Gestational Diabetes on the Gut Microbiota of Mice During Pregnancy. Indian J Microbiol 60:239–245.

21. Roselli M, Devirgiliis C, Zinno P, Guantario B, Finamore A, Rami R, Perozzi G. 2017. Impact of supplementation with a food-derived microbial community on obesity-associated inflammation and gut microbiota composition. Genes Nutr 12:25.

22. Ley RE, Turnbaugh PJ, Klein S, Gordon JI. 2006. Microbial ecology: human gut microbes associated with obesity. Nature 444:1022–3.

23. Kuang YS, Lu JH, Li SH, Li JH, Yuan MY, He JR, Chen NN, Xiao WQ, Shen SY, Qiu L, Wu YF, Hu CY, Wu YY, Li WD, Chen QZ, Deng HW, Papasian CJ, Xia HM, Qiu X. 2017. Connections between the human gut microbiome and gestational diabetes mellitus. Gigascience 6:1–12.

24. Cortez RV, Taddei CR, Sparvoli LG, Angelo AGS, Padilha M, Mattar R, Daher S. 2019. Microbiome and its relation to gestational diabetes. Endocrine 64:254–264.

25. Huang L, Thonusin C, Chattipakorn N, Chattipakorn SC. 2021. Impacts of gut microbiota on gestational diabetes mellitus: a comprehensive review. Eur J Nutr 60:2343–2360.

26. Kreznar JH, Keller MP, Traeger LL, Rabaglia ME, Schueler KL, Stapleton DS, Zhao W, Vivas EI, Yandell BS, Broman AT, Hagenbuch B, Attie AD, Rey FE. 2017. Host Genotype and Gut Microbiome Modulate Insulin Secretion and Diet-Induced Metabolic Phenotypes. Cell Rep 18:1739–1750.

27. Nardelli C, Granata I, D’Argenio V, Tramontano S, Compare D, Guarracino MR, Nardone G, Pilone V, Sacchetti L. 2020. Characterization of the duodenal mucosal microbiome in obese adult subjects by 16S rRNA sequencing. Microorganisms 8:485.

28. Kalam S, Basu A, Ahmad I, Sayyed RZ, El-Enshasy HA, Dailin DJ, Suriani NL. 2020. Recent Understanding of Soil Acidobacteria and Their Ecological Significance: A Critical Review. Front Microbiol 11:580024.

29. Chen X, Sun H, Jiang F, Shen Y, Li X, Hu X, Shen X, Wei P. 2020. Alteration of the gut microbiota associated with childhood obesity by 16S rRNA gene sequencing. PeerJ 8:e8317.

30. Dong TS, Mayer EA, Osadchiy V, Chang C, Katzka W, Lagishetty V, Gonzalez K, Kalani A, Stains J, Jacobs JP, Longo VD, Gupta A. 2020. A Distinct Brain-Gut-Microbiome Profile Exists for Females with Obesity and Food Addiction. Obesity (Silver Spring) 28:1477–1486.

31. Chen JX, Li HY, Li TT, Fu WC, Du X, Liu CH, Zhang W. 2020. Alisol A-24-acetate promotes glucose uptake via activation of AMPK in C2C12 myotubes. BMC Complement Med Ther 20:22.

32. Hu J, Lin S, Zheng B, Cheung PCK. 2018. Short-chain fatty acids in control of energy metabolism. Crit Rev Food Sci Nutr 58:1243–1249.

33. Kumar J, Rani K, Datt C. 2020. Molecular link between dietary fibre, gut microbiota and health. Mol Biol Rep 47:6229–6237.

34. He J, Zhang P, Shen L, Niu L, Tan Y, Chen L, Zhao Y, Bai L, Hao X, Li X, Zhang S, Zhu L. 2020. Short-chain fatty acids and their association with signalling pathways in inflammation, glucose and lipid metabolism. Int J Mol Sci 21:6356.

35. Pryde SE, Duncan SH, Hold GL, Stewart CS, Flint HJ. 2002. The microbiology of butyrate formation in the human colon. FEMS Microbiol Lett 217:133–9.

36. Guo J, Shao J, Yang Y, Niu X, Liao J, Zhao Q, Wang D, Li S, Hu J. 2021. Gut Microbiota in Patients with Polycystic Ovary Syndrome: a Systematic Review. Reprod Sci. doi: 10.1007/s43032-020-00430-0.

37. Guo Y, Huang ZP, Liu CQ, Qi L, Sheng Y, Zou DJ. 2018. Modulation of the gut microbiome: a systematic review of the effect of bariatric surgery. Eur J Endocrinol 178:43–56.

38. De Vadder F, Kovatcheva-Datchary P, Zitoun C, Duchampt A, Backhed F, Mithieux G. 2016. Microbiota-Produced Succinate Improves Glucose Homeostasis via Intestinal Gluconeogenesis. Cell Metab 24:151–7.

39. Nie X, Chen J, Ma X, Ni Y, Shen Y, Yu H, Panagiotou G, Bao Y. 2020. A metagenome-wide association study of gut microbiome and visceral fat accumulation. Comput Struct Biotechnol J 18:2596–2609.

40. de Moraes AC, Fernandes GR, da Silva IT, Almeida-Pititto B, Gomes EP, Pereira AD, Ferreira SR. 2017. Enterotype May Drive the Dietary-Associated Cardiometabolic Risk Factors. Front Cell Infect Microbiol 7:47.

41. Magne F, Gotteland M, Gauthier L, Zazueta A, Pesoa S, Navarrete P, Balamurugan R. 2020. The firmicutes/bacteroidetes ratio: a relevant marker of gut dysbiosis in obese patients? Nutrients 12:1474.

42. Zheng W, Xu Q, Huang W, Yan Q, Chen Y, Zhang L, Tian Z, Liu T, Yuan X, Liu C, Luo J, Guo C, Song W, Zhang L, Liang X, Qin H, Li G. 2020. Gestational diabetes mellitus is associated with reduced dynamics of gut microbiota during the first half of pregnancy. mSystems 5:e00109–e00120.

43. Wu GD, Chen J, Hoffmann C, Bittinger K, Chen YY, Keilbaugh SA, Bewtra M, Knights D, Walters WA, Knight R, Sinha R, Gilroy E, Gupta K, Baldassano R, Nessel L, Li H, Bushman FD, Lewis JD. 2011. Linking long-term dietary patterns with gut microbial enterotypes. Science 334:105–8.

44. Clarke SF, Murphy EF, O’Sullivan O, Lucey AJ, Humphreys M, Hogan A, Hayes P, O’Reilly M, Jeffery IB, Wood-Martin R, Kerins DM, Quigley E, Ross RP, O’Toole PW, Molloy MG, Falvey E, Shanahan F, Cotter PD. 2014. Exercise and associated dietary extremes impact on gut microbial diversity. Gut 63:1913–1920.

45. Conterno L, Fava F, Viola R, Tuohy KM. 2011. Obesity and the gut microbiota: does up-regulating colonic fermentation protect against obesity and metabolic disease? Genes Nutr 6:241–260.

46. Boulange CL, Neves AL, Chilloux J, Nicholson JK, Dumas ME. 2016. Impact of the gut microbiota on inflammation, obesity, and metabolic disease. Genome Med 8:42.

